# Absence of EOGT Precludes Defective Development in Fringe-null Mouse Intestine

**DOI:** 10.64898/2026.02.10.705133

**Authors:** Mohd Nauman, Pamela Stanley

**Author notes:** Dept. Pathology, Albert Einstein College of Medicine, 1300 Morris Park Ave., New York, NY, 10461.

## Abstract

Identifying biological roles for glycosyltransferases is a continuing challenge and important for defining morbidities associated with congenital disorders of glycosylation. Here we investigate the consequences to intestinal development of conditionally deleting *Lfng* alone or *Lfng, Mfng* and *Rfng* together in a mixed or *Eogt*-null genetic background. Each Fringe transfers N-acetylglucosamine (GlcNAc) to fucose (Fuc) attached to Ser or Thr by POFUT1 in a consensus sequence found in certain epithelial growth factor-like (EGF) repeats. EOGT transfers GlcNAc directly to Ser/Thr in a separate consensus sequence of the EGF repeat. Notch receptors and Notch ligands contain the largest number of EGF repeats with consensus sites for these O-glycans. Conditional deletion of *Pofut1* in mouse intestine causes similar developmental defects to deletion of *Notch1* and *Notch2* or *Dll1* and *Dll4*. LFNG also contributes to optimal Notch signaling in mouse intestine. In this work, we generated *Lfng*[F/F]:*Villin*-Cre and *Lfng*[F/F]*Mfng*[-/-]*Rfng*[-/-]:*Villin*-Cre mice in which extension of O-Fuc on EGF repeats was inhibited or prevented in intestinal epithelium. Conditional deletion of either *Lfng* alone or all three Fringe activities together led to defective intestinal development with a marked increase in goblet and Paneth cells, increased crypt width and reduced villus length. Unexpectedly, in mice globally lacking EOGT, conditional inactivation of the three Fringe genes did not lead to defective intestinal development. Thus, the absence of EOGT prevented disruption of development in Fringe-null intestine, identifying a novel role for EOGT in regulating intestinal development.

## 1. Introduction

Understanding the biological functions of mammalian glycosyltransferases is key to revealing the potential morbidities of diseases that arise from their loss or reduced activity. There are now ∼170 defined congenital diseases of glycosylation (CDG) due to mutations in genes that affect glycosylation [1] (https://www.cdghub.com/). Mutations in glycosyltransferases that transfer sugars to epidermal growth factor-like (EGF) repeats form the basis of several CDG including POFUT1-CDG [2, 3], LFNG-CDG [4], and EOGT-CDG [5]. Only about 50 proteins contain EGF repeats with consensus sequences for modification by these glycosyltransferases [6, 7]. Notch receptors and their ligands contain by far the largest number of such EGF repeats [8] (Fig. 1A). The glycosyltransferases that modify these repeats, the respective consensus sequence recognized and potential sugar extensions at each site, are summarized in Fig. 1A.

**Figure 1.**
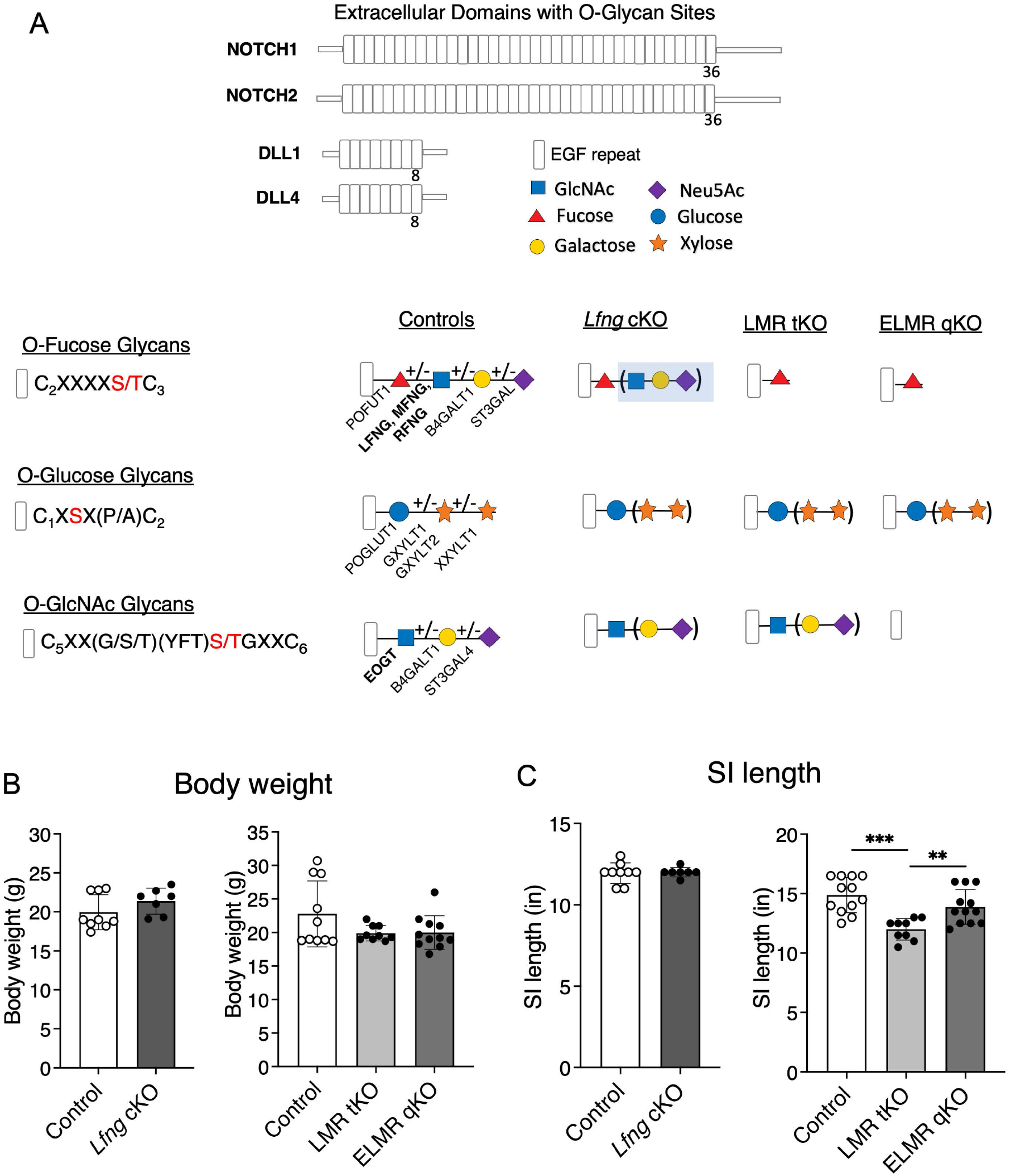
O-Glycans of EGF repeats and consequences of their removal. (A) Diagram depicting the EGF repeats of NOTCH1, NOTCH2, DLL1 and DLL4 extracellular domains. The consensus sequence recognized by the glycosyltransferase that transfers a Fuc, Glc or GlcNAc to Ser/Thr in an EGF repeat is given on the left and the sequence of sugars that may occur at each site is given on the right. The Controls column shows the full sugar extension that may occur at each site with the enzyme(s) that would be responsible for transfer [7, 25-27]. The other columns show the sequences remaining after deletion of *Lfng* alone (*Lfng* cKO) or all three Fringes together (LMR tKO) or all three Fringes in an *Eogt*-null background (ELMR qKO). Parentheses indicate potential additions. Shaded sugars in Lfng cKO reflect potential addtion by MFNG and/or RFNG. (B, C) Body weight (B) and small intestine (SI) length (C) in control and experimental mice (n ≥ 7 mice per group). *P* values from one-way ANOVA followed by Tukey’s multiple comparisons test \*\**P* < 0.01, \*\*\**P* < 0.001.

Notch signaling in intestine requires both NOTCH1 and NOTCH2 [9] stimulated by DLL1 and DLL4 [10] and has been well-studied by classical genetic approaches [11, 12]. Roles in the intestine for the glycosyltransferases that modify Notch receptors and their ligands are less well understood. Conditional deletion of *Pofut1* in intestinal epithelium, precluding the addition of O-fucose glycans to EGF repeats (Fig. 1A), markedly inhibits Notch signaling and alters intestinal cell fates [13, 14]. Global deletion of *Lfng* causes similar although milder Notch signaling defects [15]. Intestinal organoids lacking RFNG or LFNG (Fig. 1A), promote Notch signaling by modifying Notch ligands in Paneth or transit-amplifying cells, respectively [15]. MFNG is apparently not needed for Notch signaling in mouse intestine when LFNG and RFNG are present [15]. We have previously reported that global deletion of the GlcNAc-transferase EOGT (Fig. 1A) causes no apparent phenotype in intestine [14]. However, EOGT was found to promote residual Notch signaling when *Pofut1* and O-fucose glycans are absent [14].

In this paper, we characterize mouse intestine and isolated crypt cells following conditional deletion of *Lfng* by *Villin*-Cre (*Lfng* cKO) versus conditional deletion of *Lfng* in the intestine of *Mfng*.*Rfng*-null mice (LMR tKO), compared to triple Fringe gene inactivation in an *Eogt*-null background (ELMR qKO). The O-glycans expressed in intestinal epithelia of each of these genotypes are depicted in Fig. 1A. Removal of LFNG or all three Fringes in an *Eogt*[+/-] heterozygous background led to altered cell fates and disrupted intestinal development. However, when the three Fringe genes were deleted in mice globally lacking EOGT, there was no apparent effect on cell fate decisions and intestinal development appeared normal in ELMR qKO intestine. Therefore, the absence of EOGT protected intestinal epithelium from the cell fate changes that arose when all Fringes were removed. This unexpected result highlights the complex roles that glycosyltransferases play in the regulation of intestinal development and identifies a new functional role for EOGT in the intestine.

## 2. Materials and Methods

### 2.1. Mice

Mice harboring a targeted inactivating mutation in the *Eogt* gene were generated at Nagoya University as previously described [16]. *Lfng*[F/F] mice were a generous gift from Dr. Sean Egan (Hospital for Sick Children Research Institute, Toronto, Ontario, Canada) [17]. *Mfng*[-/-]*Rfng*[-/-] double KO mice were kindly provided by Dr. Susan E. Cole (The Ohio State University, Columbus, OH, USA) [18]. Mice expressing *Villin*-Cre (B6.Cg-Tg(Vil1-cre)1000Gum/J, JAX strain #021504) [19] were originally procured from The Jackson Laboratories, Portland, MA and used previously [14]. All mice were backcrossed extensively to C57BL/6J.. Experimental groups were generated as follows: *Lfng*[F/F] mice were crossed to *Villin*-Cre heterozygotes to obtain *Lfng*[F/F] mice with one copy of the *Villin*-Cre transgene (*Lfng* cKO) and no *Villin*-Cre (Control); to obtain LMR tKO mice *Eogt*[+/-]*Lfng*[F/F]*Mfng*[-/-]*Rfng*[-/-]:*Villin*-Cre mice were generated and to obtain ELMR qKO mice *Eogt*[-/-]*Lfng*[F/F]*Mfng*[-/-]*Rfng*[-/-]:*Villin*-Cre mice were generated by crossing *Eogt* null, *Lfng*[F/F], *Mfng*[-/-]*Rfng*[-/-] and *Villin*-Cre heterozygous mice in appropriate combinations. *Eogt*[+/-]*Lfng*[F/+]*Mfng*[+/-]*Rfng*[+/-] and *Eogt*[+/-]*Lfng*[F/+]*Mfng*[+/+]*Rfng*[+/-] mice were used as controls for LMR tKO and ELMR qKO mice. PCR of genomic DNA (gDNA) was used to genotype mice with the primer sets in Supplementary Table S1.

To obtain intestinal tissue, mice of both genders at 6-9 weeks of age were euthanized by CO_2_ asphyxiation followed by cervical dislocation. Following euthanasia, body weight was determined, small intestine was dissected, the length measured, and tissue samples were processed as described below. All procedures were approved by the Albert Einstein Institutional Animal Care and Use Committee (IACUC) under protocol numbers 20170709 and 00001311. Experiments were carried out in compliance with the approved protocols and relevant regulations.

### 2.2. Histopathology

The jejunum was fixed in 10% neutral buffered formalin for 48 hr before processing through a graded ethanol series for paraffin embedding. Sections of 5 μm were stained using hematoxylin and eosin (H&E) or Alcian Blue. Reagents were purchased from Sigma (St. Louis, MO, USA). Slides were scanned with a 3D Histech P250 High-Capacity Slide Scanner. Morphometric analyses including villus length, crypt depth, crypt width, and crypt length were performed using CaseViewer 2.4 software. Paneth cells and goblet cells were counted manually in scanned slides.

### 2.3. Isolation of crypts and villi

Crypt and villus fractions were isolated from small intestine after washing with cold phosphate-buffered saline containing 1 mM CaCl_2_, 1 mM MgCl_2_, and 1 mM dithiothreitol (PBS Ca/Mg/DTT). The intestine was cut into four segments, extensively flushed with the same buffer, one end tied, then everted using a gavage needle, and filled about 75% with PBS Ca/Mg/DTT before being tied off at the other end. Intestinal tubes were incubated in buffer A (96 mM NaCl, 1.5 mM KCl, 27 mM Na-Citrate, 8 mM KH_2_PO_4_, 5.6 mM Na_2_HPO_4_, and 1 mM DTT) at 37°C with continuous shaking for 10 min. Subsequently, the tissue was incubated 4 sequential times in buffer B (1.5 mM EDTA, 0.5 mM DTT, 0.1% BSA) with shaking at 37 °C for 10, 10, 30, and 20 min, respectively. Each collection of buffer B was centrifuged at 4ºC to isolate intestinal fractions I through IV. The relative enrichment of villus and crypt populations in these fractions was assessed by qRT-PCR analysis of marker genes (Supplementary Fig. S1). Fractions were snap-frozen in liquid nitrogen and stored at −80°C. Deletion of *Lfng* via *Villin-Cre* in crypts was validated by PCR genotyping of gDNA (Supplementary Fig. S2).

### 2.4. Flow cytometry

Everted small intestine tubes were gently scraped to remove villi, placed into 20 ml of cold PBS (Ca^2+^/Mg^2+^-free) supplemented with 2 mM EDTA and 2 mM glutamine, and held on ice for 20 min. The tubes were then moved to 20 ml cold PBS (Ca^2+^/Mg^2+^-free) containing 2 mM glutamine (PBS/glutamine) and manually agitated for 30 sec. The washings from 4 additional 30 sec washing steps in fresh PBS/glutamine were combined and passed through a 70 μm strainer into a 50 ml Falcon centiruge tube precoated with 1% bovine serum albumin (BSA fraction V; GeminiBio, West Sacramento, CA) in PBS (Ca^2+^/Mg^2+^ free). The filtrate was centrifuged at 1500 rpm for 10 min at 4°C to collect crypts. Crypts were dissociated into single cells using 3 mL enzyme-free dissociation buffer (Gibco/Thermo Fisher Scientific, Waltham, MA) in a 37°C water bath for 10 min, with gentle pipetting every 30 sec. Following the addition of 7 ml alpha-MEM (Gibco), cells were passed through a 40 μm strainer, centrifuged, and resuspended in Zombie NIR dye (1:7000 dilution in PBS (Ca^2+^/Mg^2+^-free); BioLegend). After rocking for 30 min at 4°C, cells were washed in PBS (Ca^2+^/Mg^2+^-free), fixed in 4% paraformaldehyde (PFA; Electron Microscopy Sciences, Hatfield, PA) in PBS (Ca^2+^/Mg^2+^-free) for 15 min at 4°C with rocking, followed by 3 washes in cold FACS binding buffer (FBB; Hank’s balanced salt solution (Millipore Sigma Corning, Burlington, MA), 2% BSA, 1 mM CaCl_2_ and 0.1% sodium azide), and stored in FBB at 4°C. Samples were analyzed by flow cytometry within 1-6 days of preparation. Cells were assessed for NOTCH1 surface expression with anti-NOTCH1 extracellular domain (ECD) antibody (Ab AF5267; R&D Systems, Inc., Minneapolis, MN) and binding of Notch ligands DLL1-Fc (R&D Systems, Inc.) and DLL4-Fc (Acro Biosystems, Newark, DE).

Flow cytometry was performed on 6 different days each time with crypt cells from one Control, one LMR tKO and one ELMR qKO intestine to analyze cell surface NOTCH1 expression, and DLL1-Fc and DLL4-Fc binding. Approximately 10^6^ crypt cells per tube were incubated in CD16/CD32 Fc blocker (1:40 in FBB) for 15 min on ice. Cells were then stained for 30 min on ice with antibodies in FBB against CD45 (1:400), CD44 (1:800), CD24 (1:800) plus either anti-NOTCH1 ECD or 2 μg soluble NOTCH ligand (DLL1-Fc or DLL4-Fc) in Fc-block solution. Sources of antibodies are given in Supplementary Table S2. After three washes in FBB, detection of bound Ab or ligand used rhodamine Red-X-conjugated donkey anti-sheep IgG (1:600) or Fc-specific anti-IgG-DyLight 405 (1:100) in FBB for 30 min on ice. Background staining was established using cells incubated without Ab to NOTCH1 ECD or Ab to Fc. Cells were washed three times in FBB, resuspended in 300 μL FBB, and filtered into flow cytometry tubes. 300,000 events were collected using a Cytek Aurora flow cytometer, and data was analyzed using FlowJo software (BD Biosciences). Mean fluorescence index (MFI) was determined for anti-NOTCH1 ECD, DLL1-Fc and DLL4-Fc binding in each sample. MFI for Control mice was used to calculate the relative binding of anti-NOTCH1 Ab and anti-Fc Ab.

### 2.5. Quantitative RT-PCR (qRT-PCR)

Total RNA was isolated from ∼10^7^ frozen crypts (Fraction IV) or villi (Fraction I) obtained as described above, using TRIzol reagent (Thermo Fisher Scientific, Waltham, MA). RNA was resuspended in RNase-free water, and 1 µg RNA quantified by Nanodrop ND1000 Spectrophotometer (Thermo Fisher Scientific), was used to synthesize cDNA in 20 µl using the Verso cDNA synthesis kit (Thermo Fisher Scientific). For amplification, cDNA was combined with PowerUp SYBR Green master mix (Thermo Fisher Scientific) and primers at a final concentration of 750 nM each. Quantitative RT-PCR was performed in the Vii7 Real-Time PCR system (Applied Biosystems, Foster City, CA) with a 40-cycle program. Experiments were conducted in triplicate in a 384-well plate format. Relative gene expression levels were determined using the log2^ddCT^ method [20, 21] normalized to the ubiquitously-expressed genes *Gapdh* and *Hprt*. Sequences of primers are given in Supplementary Table S3.

## 3. Results

### 3.1. Conditional deletion of Lfng, Mfng and Rfng in the intestine

To determine effects on intestinal development of Fringe glycosyltransferases that modify Notch receptors and Notch ligands, conditional deletion in intestinal epithelium by *Villin*-Cre was performed on *Lfng* in various mutant backgrounds as described in Methods. The predicted structure of O-glycans on EGF repeats in the different mutants is summarized in Fig. 1A. We previously found that deletion of *Pofut1* using *Villin*-Cre causes severe disruption of Notch signaling and intestinal development in mice [13, 14]. By contrast, global deletion of *Eogt* had no apparent effect on cell fate decisions in the intestine [14]. However, conditional deletion of *Pofut1* in an *Eogt*-null background revealed that EOGT supports Notch signaling in the absence of POFUT1 [14]. To investigate additional interactions between glycosyltransferases in intestinal development, we generated mouse models to compared deletion of *Lfng* in intestine (*Lfng* cKO), versus deletion of *Lfng* in the intestine of mice lacking MFNG and RFNG (LMR tKO) versus deletion of *Lfng* in the intestine of mice lacking EOGT, MFNG and RFNG (ELMR qKO) as described in Methods.

No significant differences were observed in body weight or small intestine length between *Lfng* cKO intestine and controls (Fig. 1B and C). Similarly, body weight did not differ between LMR tKO or ELMR qKO mice compared to their respective controls (Fig. 1B). By contrast, small intestine length was significantly reduced in LMR tKO mice (Fig. 1C). Interestingly, the absence of *Eogt* in ELMR qKO mice precluded the reduction in small intestine length following LMR tKO so that small intestine length was similar to controls in ELMR qKO intestine (Fig. 1C).

### 3.2. Goblet and Paneth cells in Lfng cKO, LMR tKO and ELMR qKO intestine

Increased numbers of secretory cells is a hallmark of reduced Notch signaling in the small intestine [9, 10, 13, 14]. A major secretory cell arising from *Lgr5*+ stem cells is the goblet cell. The number of goblet cells in the crypts of *Lfng* cKO intestine did not increase significantly (Fig. 2A and C) but goblet cells in villi of *Lfng* cKO mice were increased compared to controls (Fig. 2B and C). By contrast, goblet cell numbers were elevated in crypts of LMR tKO mice but remained unchanged in villi (Fig. 2A-C). This difference indicated an effect of MFNG and/or RFNG on cell fate decisions in LMR tKO intestine. The absence of *Eogt* in ELMR qKO mice prevented the increase in goblet cell numbers observed in LMR tKO intestine. Goblet cell numbers remained at control levels in crypts and villi of ELMR qKO intestine (Fig. 2A-C).

**Figure 2.**
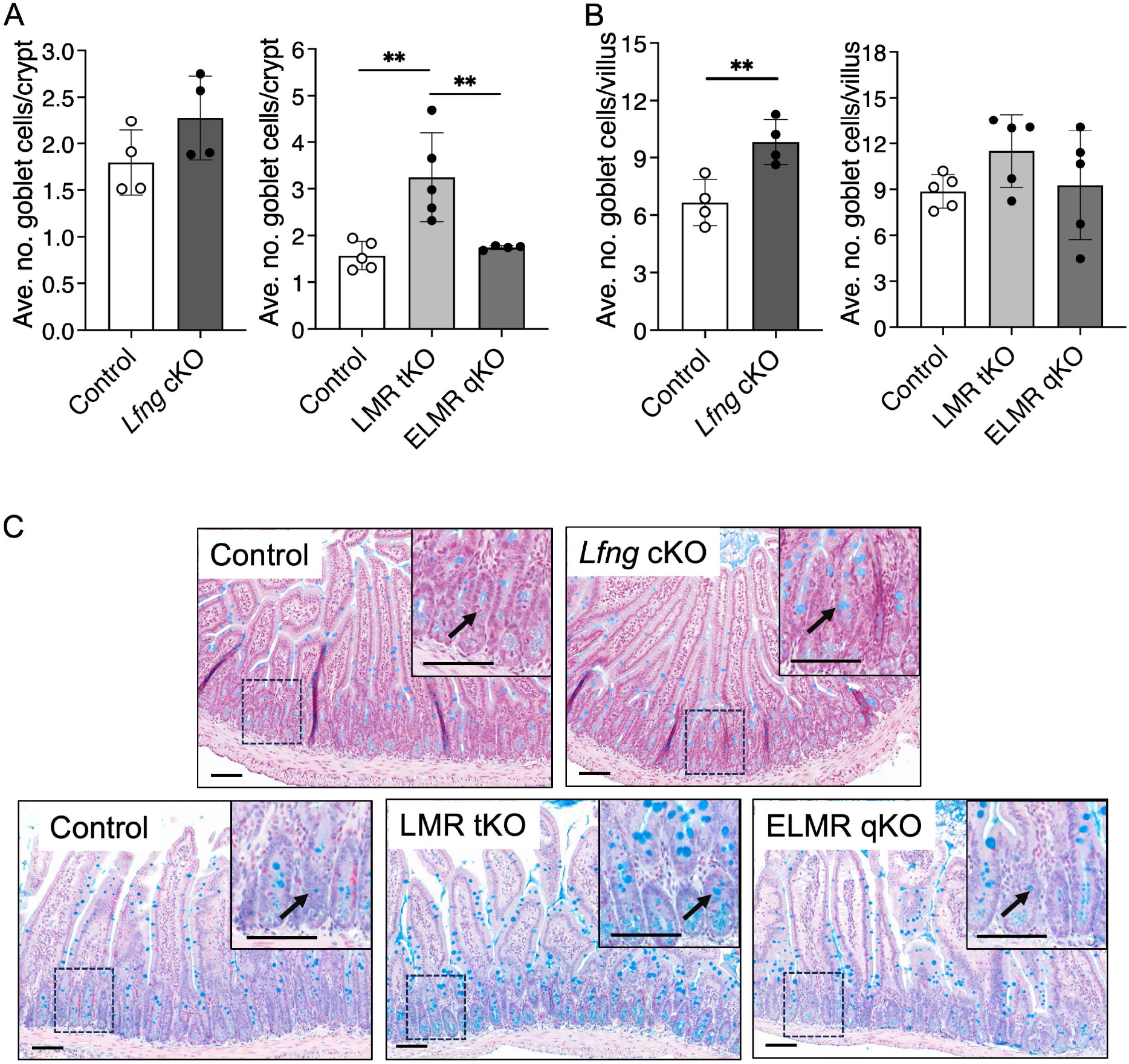
Goblet cells in *Lfng* cKO, LMR tKO and ELMR qKO intestine. (A, B) Number of goblet cells in the crypts (A) and villi (B) of experimental and control intestine (50 crypts or 35 villi were analyzed in n ≥ 4 mice per group). *P* values from unpaired, two-tailed Student’s t test \*\**P* < 0.01 (Control vs *Lfng* cKO) or one-way ANOVA followed by Tukey’s multiple comparisons test \*\**P* < 0.01 (Control vs LMR tKO vs ELMR qKO). (C) Representative images showing goblet cells (arrowheads). Scale bar 100 μm.

Paneth cells are another type of secretory cell that originate from intestinal *Lgr5*+ cells and reside within the base of crypts. Conditional deletion of *Lfng* led to increased Paneth cell numbers in crypts (Fig. 3A and C). Paneth cell numbers were similarly elevated in the crypts of LMR tKO intestine but were not increased when inactivation of the three Fringe genes occurred in an *Eogt-*null background in ELMR qKO intestine (Fig. 3B and C).

**Figure 3.**
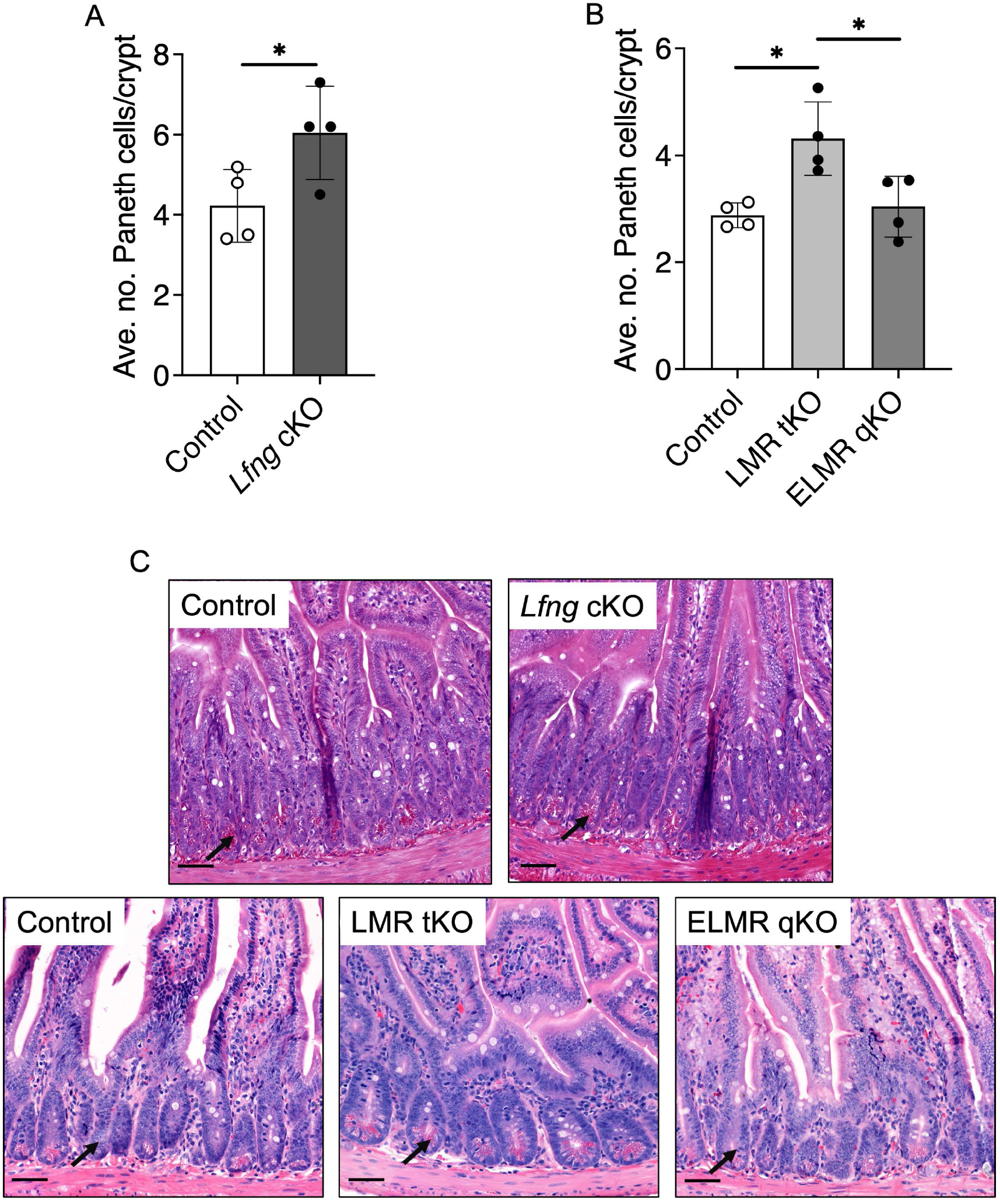
Paneth cells in *Lfng* cKO, LMR tKO and ELMR qKO intestine. (A,B) Number of Paneth cells in the crypts of experimental mice (50 crypts were analyzed in n ≥ 4 mice per group). *P* values from unpaired, two-tailed Student’s t test \**P* < 0.05 (Control vs *Lfng* cKO) or one-way ANOVA followed by Tukey’s multiple comparisons test \*\**P* < 0.01 (Control vs LMR tKO vs ELMR qKO). (C) Representative images showing Paneth cells in crypts (arrowheads). Scale bar 50 μm.

### 3.4. Further changes in LMR tKO and ELMR qKO intestine

Intestinal homeostasis maintains the balance between absorptive enterocytes (primarily villus cells) and secretory lineages (primarily resident in crypts). Reduced Notch signaling disrupts cell fate differentiation increasing secretory cell production at the expense of enterocytes leading to shortened villi. Secretory cell accumulation in crypts leads to crypt expansion leading to increased depth and width. *Lfng* cKO intestines exhibited no differences in villus length or crypt width from controls (Fig. 4A and B). However, crypt depth was increased in *Lfng* cKO intestine (Fig. 4C). In LMR tKO intestine, villus length was significantly reduced consistent with decreased formation of absorptive enterocytes (Fig. 4A). Crypt width was increased in LMR tKO intestine (Fig. 4B) but there was no change in crypt depth (Fig. 4C). In the absence of EOGT, the loss of the three Fringes in *Eogt*-null intestine did not cause any changes in villus length, crypt width or crypt depth in ELMR qKO intestine (Fig. 4A-C).

**Figure 4.**
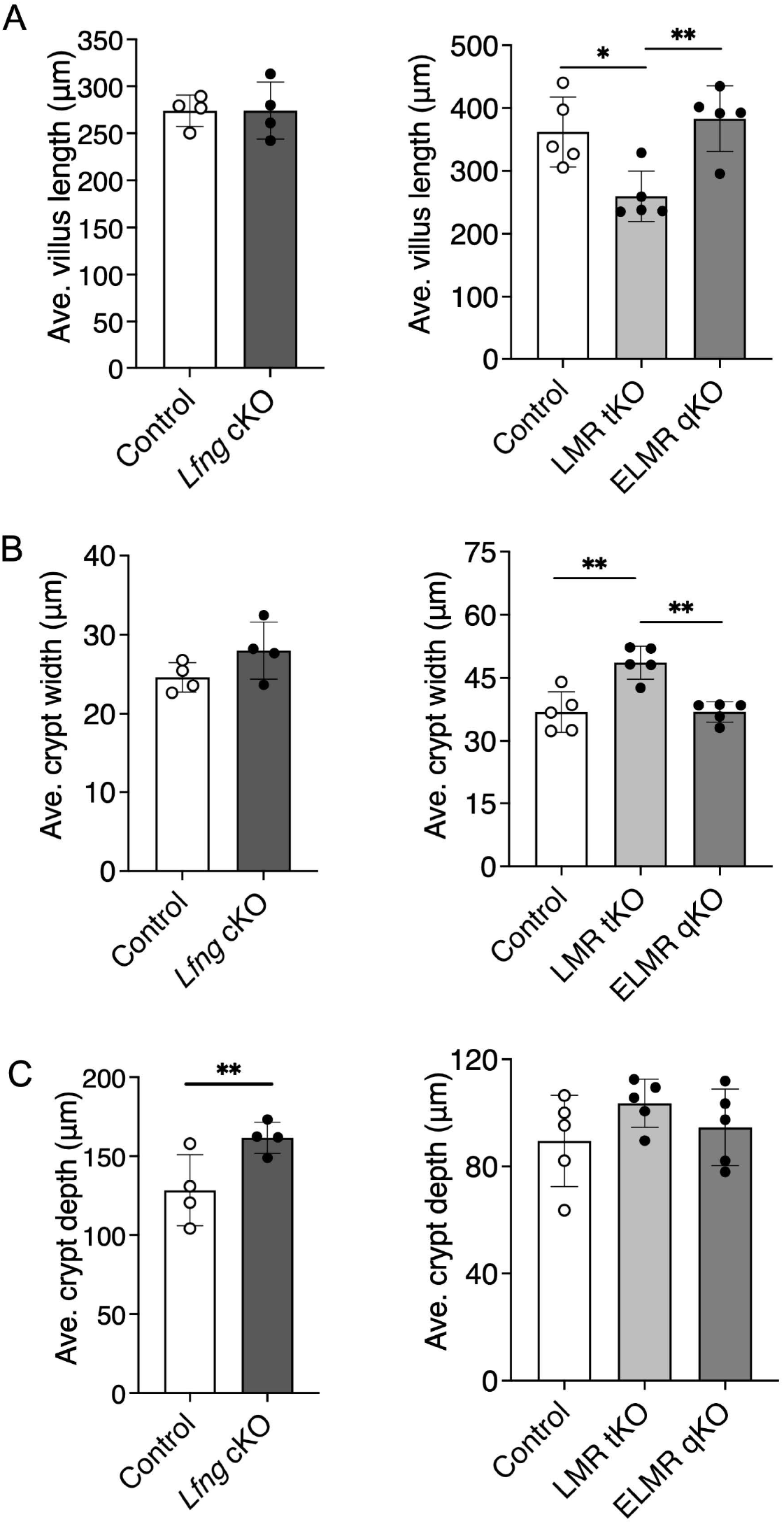
Morphometry of crypts and villi in *Lfng* cKO, LMR tKO and ELMR qKO intestine. (A) villus length (20 villi were analyzed in n ≥ 4 mice per group), (B) crypt width and (C) crypt depth (50 crypts were analyzed in n ≥ 4 mice per group). *P* values from unpaired, two-tailed Student’s t test **P < 0.01 (Control vs *Lfng* cKO) or one-way ANOVA followed by Tukey’s multiple comparisons test \**P* < 0.5, \*\**P* < 0.01 (Control vs LMR tKO vs ELMR qKO). Scale bar 100 μm.

### 3.5. Notch ligand binding to LMR tKO and ELMR qKO intestinal stem cells

To investigate Notch ligand binding to Notch receptors in intestinal stem cells (ISC), single cell suspensions were prepared from isolated crypts and analyzed by flow cytometry. Viable, single cells were gated on CD44+CD45−CD24− ISC (Supplementary Fig. S3). Cell surface expression of NOTCH1 was determined using Ab to the ECD, and binding of DLL1-Fc or DLL4-FC was determined using anti-Fc Ab. Six experiments were performed, each including ISC prepared from crypts of 3 mice - control, LMR tKO and ELMR qKO. Representative flow cytometry profiles and histograms of mean fluorescence index (MFI) for NOTCH1, DLL1-Fc and DLL4-Fc are shown in Fig. 5A-C. MFI data for mutant ISC was normalized to control ISC in each experiment. The expression of NOTCH1 on ISC cell surface was quite variable but did not differ significantly between groups (Fig. 5D). Binding of DLL1-Fc and DLL4-Fc also did not differ significantly from control in either LMR tKO or ELMR qKO ISC (Fig. 5D). Therefore, Notch receptors carrying EGF repeats with Fuc that was not extended by GlcNAc in LMR tKO ISC, and Notch receptors carrying unextended Fuc and no O-GlcNAc glycans in ELMR qKO ISC, were expressed at the cell surface and bound Notch ligands.

**Figure 5.**
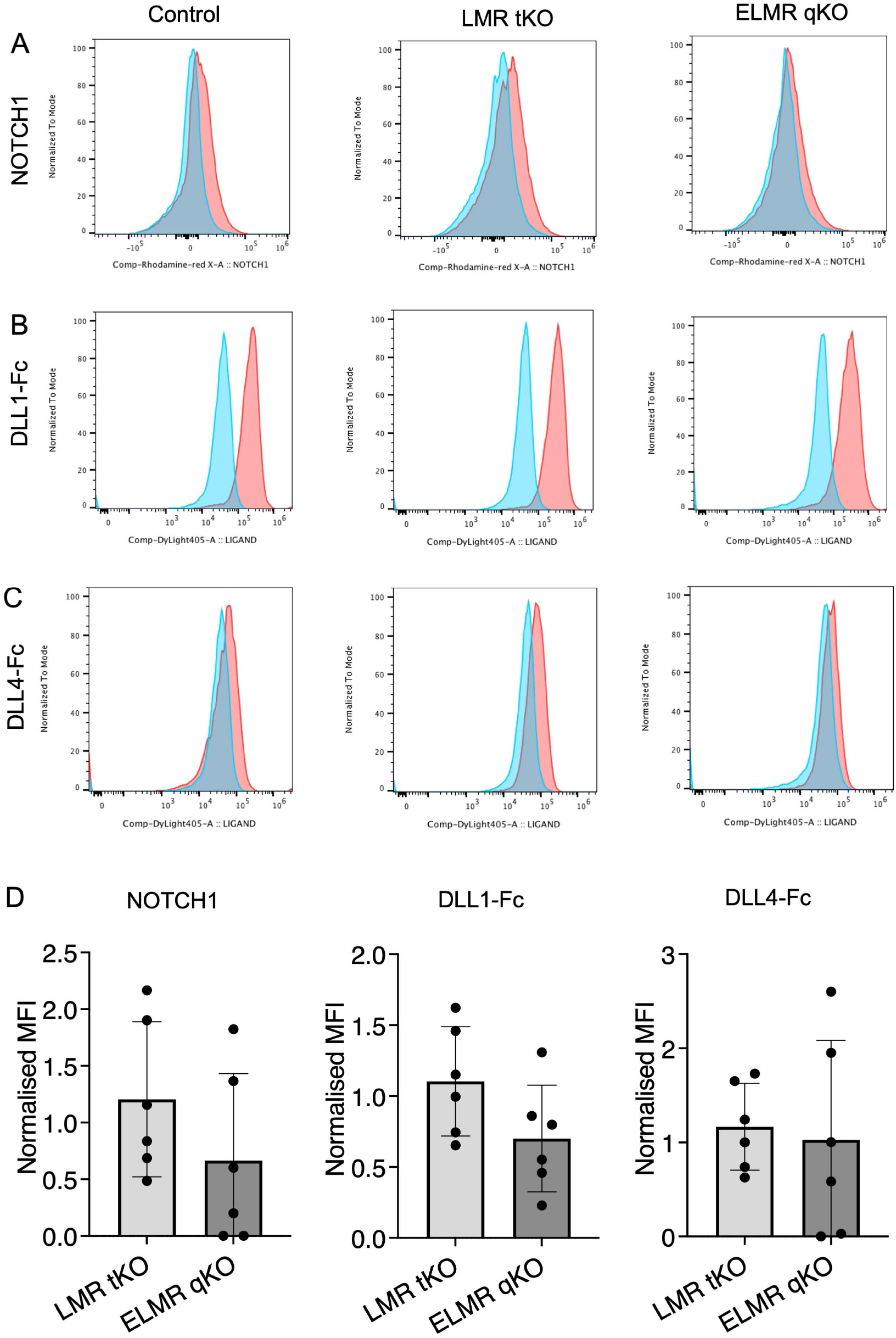
Notch ligand binding to ISC in crypts. (A-C) Flow cytometry profiles of anti-NOTCH1 ECD Ab, DLL1-Fc and DLL4-Fc binding to ISC from Control, LMR tKO and ELMR qKO crypts. Pink profiles show Ab to NOTCH1 ECD or Notch ligand-Fc binding, blue profiles show 2° antibody non-specific binding. (D) MFI for Ab to NOTCH1 ECD, DLL1-Fc and DLL4-Fc binding to LMR tKO and ELMR qKO normalized to MFI of Control in each of six experiments performed over 6 days. MFI values varied between experiments so each experiment was analyzed separately. One-way ANOVA followed by Tukey’s multiple comparisons test was used to determine *P* values.

### 3.6. Gene expression in Lfng cKO, LMR tKO and ELMR qKO crypts

To assess gene expression changes in crypts isolated from *Lfng* cKO, LMR tKO and ELMR qKO intestine, expression of Notch signaling target genes, intestinal lineage marker genes and Notch pathway genes were quantitated by qRT-PCR. Fractionation of intestinal tube washings yielded comparable crypt versus villus enrichment across all genotypes (Supplementary Fig. S1A and B). Expression of *Hes7*, a Notch target gene, was increased in *Lfng* cKO and LMR tKO mice but did not change from control levels in ELMR qKO crypts (Fig. 6A). *Lyz1*, a marker for Paneth cells, was elevated in *Lfng* cKO but not in LMR tKO crypts (Fig. 6B). Chromogranin A, a marker of secretory enteroendocrine cells, was increased in LMR tKO crypts but was unchanged from control levels in ELMR qKO crypts (Fig. 6B). No significant changes from controls were observed in stem cell markers across *Lfng* cKO, LMR tKO, and ELMR qKO crypts (Fig. 6C). Interestingly, expression of *Notch1, Dll1*, and *Dll4* was increased in *Lfng* cKO but not in LMR tKO crypts (Fig. 6D). Interpretation of these expression data is discussed below.

**Figure 6.**
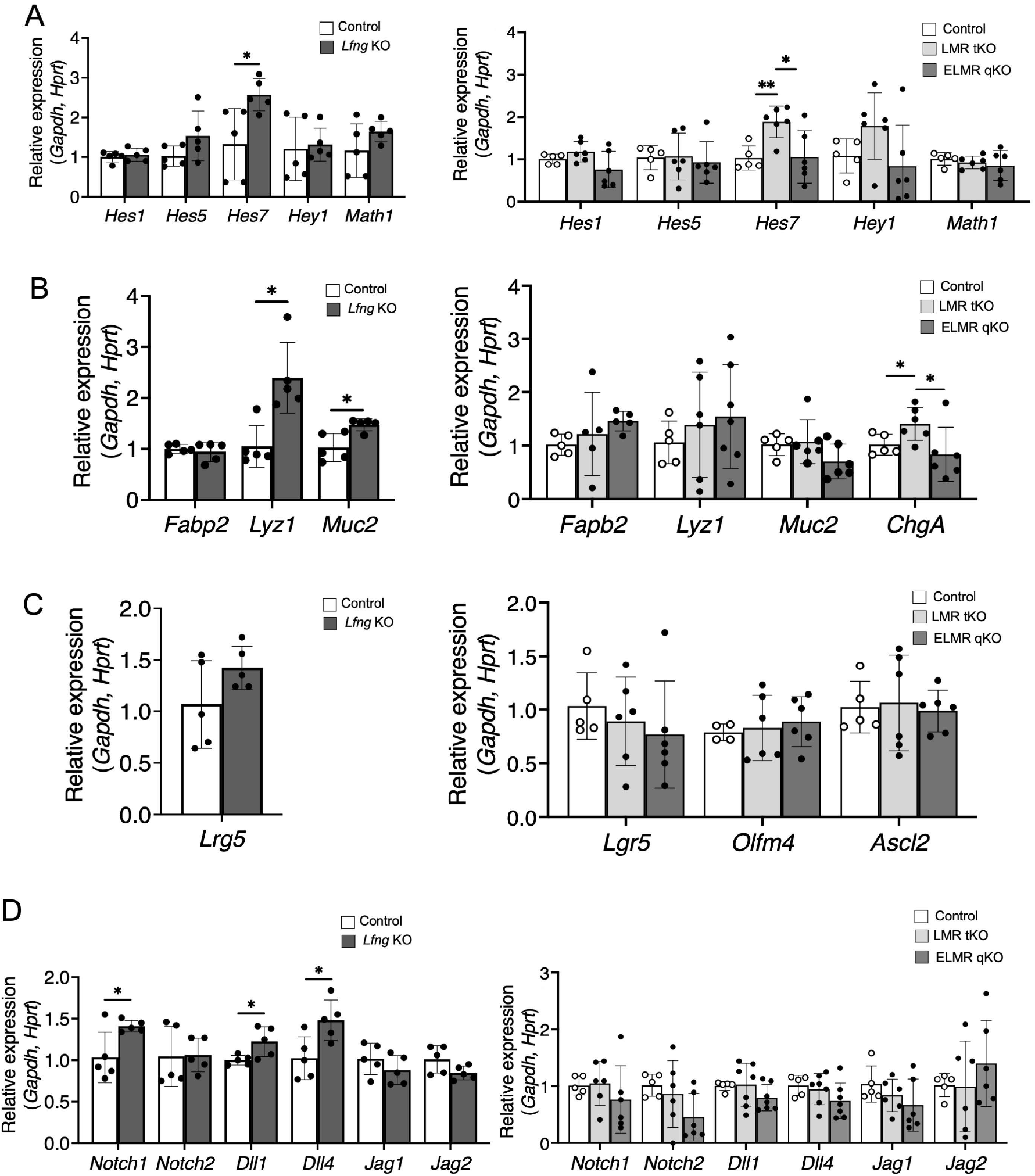
Gene expression in intestinal crypts. (A-D) qRT-PCR was performed on cDNA prepared from intestinal crypts. (A) Expression of key Notch signaling target genes in crypts of experimental mice. (B) Expresson of marker genes expressed in different intestinal cell lineages. (C) Expression of marker genes in ISC. (D) Expression of Notch receptors and Notch ligands. (n ≥ 5 mice per group). *P* values were determined by unpaired, two-tailed Student’s t test \**P* < 0.5, \*\**P* < 0.01. Additionally, statistical analysis between Control, LMR tKO and ELMR qKO was tested by One-way ANOVA followed by Tukey’s multiple comparisons test. By that analysis, *Hes7* expression increased in LMR tKO compared to Control (\**P* < 0.5) and decreased in ELMR qKO compared to LMR tKO (\**P* < 0.5).

## 4. Discussion

Here we report the unexpected finding that, in the absence of EOGT, deletion of all three Fringe genes in mouse intestine has no apparent consequences, whereas in the presence of EOGT, cell fates and development in intestine are markedly disrupted by the removal of all Fringe activity. Because EOGT and the Fringes are all GlcNAc-transferases known to modify Notch receptors and their ligands on EGF repeats, and because the developmental disruption in LMR tKO intestine is similar to (though milder than) that observed when NOTCH1 and NOTCH2 or DLL1 and DLL4 or POFUT1 are deleted, it is tempting to interpret our findings in the context of Notch signaling regulated by O-glycans [22]. The O-fucose glycans transferred to NOTCH1 EGF repeats by POFUT1 and extended by LFNG/MFNG/RFNG are important for DLL1 binding [23]. DLL1-Fc binding is reduced in *Pofut1*[-/-] ISC and further reduced in *Eogt*[-/-]*Pofut1*[-/-] ISC [14], consistent with inhibition of Notch signaling due to reduced Notch ligand binding. From an O-glycan perspective our findings predict that when EOGT is present and O-GlcNAc glycans are added to EGF repeats, those glycans facilitate or promote the disrupted development observed in LMR tKO intestine, but when EOGT is absent and there are no O-GlcNAc glycans on EGF repeats, the additional removal of all Fringes that extend O-Fuc with GlcNAc does not lead to disrupted development. Understanding mechanistic bases of these findings will be the focus of future work. The trivial explanation that LMR tKO cells were excluded from crypts when EOGT was absent was found not to be the case. PCR genotyping of crypt gDNA showed the same signal intensity for deleted *Lfng* allele in LMR tKO and ELMR qKO cypts (Supplementary Fig. S2).

Notch signaling was investigated in our mutant models by characterizing Notch ligand binding and gene expression in fractioned crypts. No significant reduction in NOTCH1 ECD Ab, DLL1-Fc or DLL4-Fc binding to LMR tKO or ELMR qKO ISC was observed (Fig. 5). Therefore, NOTCH1 cell surface expression was similar in LMR tKO and ELMR qKO ISC. Gene expression changes were few in LMR tKO and ELMR qKO crypts and did not suggest reduced expression of typical Notch target genes (Fig. 6). For example, deletion of *Pofut1* in intestine causes reduced expression of Notch signaling target *Hes1* and increased expression of *Math1/Atoh1* [13, 14] and *Hes1* expression is further reduced in *Eogt*[-/-]*Pofut1*[-/-] crypts [14]. By contrast, there was no significant change in *Hes1* or *Math1/Atoh1* expression in *Lfng* cKO, LMR tKO or ELMR qKO crypts (Fig. 6). However, global KO of *Lfng* was previously reported to cause reduced *Hes1* and increased *Atoh1* expression in ISC [15]. The differences observed between KO and cKO of *Lfng* may be due to genetic background or the loss of all LFNG activity in global *Lfng* KO verus loss only in intestinal epithelium in *Lfng* cKO. There was a significant increase in *Hes7* expression in *Lfng* cKO as well as in LMR tKO crypts which did not occur in ELMR qKO crypts, consistent with the LMR tKO phenotype not manifesting in an *Eogt*-null background. *Hes7* is a known target gene positively regulated by Notch signaling [24]. The fact that *Hes7* expression went up in *Lfng* cKO and LMR tKO crypts indicates that Notch signaling was increased. Consistent with this was an increase in expression of *Notch1, Dll1* and *Dll4* in *Lfng* cKO crypts. However, these increases were not observed in LMR tKO crypts suggesting an influence of the presence of MFNG and/or RFNG in promoting the *Lfng* cKO crypt phenotype.

In the final analysis, while the Paneth and goblet secretory cell number increases, and the crypt and villus changes in *Lfng* cKO and LMR tKO intestine mimic the developmental phenotype expected from reduced Notch signaling and observed previously for *Lfng*[-/-] intestine [15], the fact that gene expression changes were few and opposite to expectations in *Lfng* cKO intestine requires a more complicated interpretation. A complicating factor is the complex nature of intestinal crypts. Further purification of different cell types will allow more definitive distinctions in gene expression changes. Organoid co-culture experiments with purified wild-type and mutant cells will allow functional roles of Fringes in different cell types to be identified.

## Acknowledgements

We thank Mohammad Asad and Subha Sundaram for technical assistance.

## CRediT authorship contribution statement

**Mohd Nauman:** Writing – Original draft, Investigation, Methodology, Formal analysis, Data curation **Pamela Stanley:** Writing – review & editing, Supervision, Conceptualization, Data curation, Project administration, Funding acquisition.

