## Supplementary data for "Absence of EOGT Precludes Defective Development in Fringe-null Mouse Intestine"

**Table of Contents**

**Supplementary Tables**

| | |
|---|---|
| Supplementary Table S1 | Primer sequences used for genotyping |
| Supplementary Table S2 | Antibodies used in flow cytometry |
| Supplementary Table S3 | Primer sequences used in qRT-PCR |

**Supplementary Figures**

| | |
|---|---|
| Supplementary Figure S1 | Validation of villus and crypt fractions |
| Supplementary Figure S2 | Validation of *Lfng* deletion by *Villin1*-Cre in gDNA isolated from crypts of LMR tKO and ELMR qKO intestine |
| Supplementary Figure S3 | Gating strategy for flow cytometry analysis |

Supplementary Table S1. Primer sequences used for genotyping

| Gene | Allele | Sequence (5’→3)’ |
|---|---|---|
| *Eogt* | *Wild type*<br>*Wild type*<br>*Knockout* | *F: CCACCCGACCCCTGCCAGAACATAATGCTCTCTTGCATC*<br>*R: GCTGTCGCCAGAGGAGAGAGTGGGTGCTTACTTAC*<br>*R: CCAAGGCGGTCTTGGCCCAT* |
| *Rfng* | *Wild type*<br>*Wild type*<br>*Knockout* | *F: GCAATTGGTGTTGATCATGC*<br>*R: CGTTTTCTCCCACAGACGTT*<br>*R: TTCAACATCAGCCGCTACAG* |
| *Mfng* | *Wild type*<br>*Wild type*<br>*Knockout* | *F: AGACCCAGCTTCCTCCTTTC*<br>*R:CCGACTCCCCAATCAGTCT*<br>*R: GCATGCCTAGCCCTGTGTAT* |
| *Lfng* | *Flox*<br>*Flox* | *F: ACCTGTCTGAAGTTGGAGAGTGAG*<br>*R: AGCAGTTGGTGAGCACCACATTGC* |
| *Vil1* | *Wild type*<br>*Wild-type*<br>*Villin1-Cre* | *F: GCCTTCTCCTCTAGGCTCGT*<br>*R: TATAGGGCAGAGCTGGAGGA*<br>*R: AGGCAAATTTTGGTGTACGG* |

Supplementary Table S2. Antibodies used in flow cytometry

| Antigen | Fluorochrome | Isotype | Clone | Cat. no. | Company |
|---|---|---|---|---|---|
| NOTCH1 aa19-526 | - | Ag-purified sheep IgG | Polyclonal | AF5267 | R&D Systems Inc., Minneapolis, MN |
| CD45 | PerCP | Rabbit IgG2b, κ | 30-F11 | 103129 | Biolegend, San Diego, CA |
| CD44 | PE-Cyanine 7 | Rabbit IgG2b, κ | IM7 | 25-0441-81 | eBioscience, San Diego, CA |
| CD24 | BV510 | Rabbit IgG2b, κ | M1/69 | 101831 | Biolegend, San Diego, CA |
| Anti-sheep IgG | Rhodamine Red-X | Donkey anti-sheep IgG | Polyclonal | 713-295-147 or 713-295-003 | Jackson ImmunoResearch, Lab. Inc., West Grove, PA |
| Anti-human Fcγ | Dylight™ 405 | Goat anti-human IgG | Polyclonal | 109-476-170 | Jackson ImmunoResearch, Lab. Inc., West Grove, PA |

Supplementary Table S3. Primer sequences used in qRT-PCR

| Gene | Sequence (5’→3)’ |
|---|---|
| *Hes1* | *F: AGCTGGAGAGGCTGCCAAGGTTT*<br>*R: ACATGGAGTCCGAAGTGAGCGAG* |
| *Hes5* | *F: GGACCAGAGGATGAGCTCGTT*<br>*R: AGGAGGGAGCCTTCGGAAGA* |
| *Hes7* | *F: GAGCGAGCTGAGAATAGGGA*<br>*R: GGCTTCGCTCCCTCAAGTAG* |
| *Hey1* | *F: TGAGCTGAGA AGGCTGGTAC*<br>*R: ACCCCAAACTCCGATAGTC* |
| *Math1* | *F: ATGCACGGGCTGAACCA*<br>*R: TCGTTGTTGAAGGACGGGATA* |
| *Notch1* | *F: GAGATGCCCTCGGACCAATC*<br>*R: GGCGAAAGCGAGAACACA* |
| *Notch2* | *F: TGTACCAGATCCCAGAGATGC*<br>*R: GTCAGATGCAGAGTGTGGTGA* |
| *Dll1* | *F: GTCTGCCAGGGTGTGATGAC*<br>*R: CGGATGCACTCATCGCAGTA* |
| *Dll4* | *F: AGTGCCAGAACAGAGGTCCAA*<br>*R: CAGGGACTTCGGGCACAAT* |
| *Jag1* | *F: TGACATGGATAAACACCAGCA*<br>*R: GCAGCCCACTGTCTGCTATAC* |
| *Jag2* | *F: ATTGTAGCAAGGTATGGTGCG* |

| | |
|---|---|
| | *R: GCACAGTTGTTGTCCAAATGA* |
| *Lgr5* | *F: TCTCCTACATCGCCTCTGCT*<br><br>*R: TTCCTCCGGAACCTGTCTCA* |
| *Olfm4* | *F: ATTCGCTATGGCCAAGGAGG*<br><br>*R: GAGGGGCCGATTCACATCAA* |
| *ChgA* | *F: GAAGTGCG CCTGGAAGTCA*<br><br>*R: GATCCTCTCGTCTCCTTGGA* |
| *Fabp2* | *F: CTAGAGACACACACAGCTGAGATCATGG*<br><br>*R: GCAATCAGCTCCTTTCCATTGTCTACAC* |
| *Hprt* | *F: TCAGTCAACGGGGGACATAAA*<br><br>*R: GGGGCTGTACTGCTTAACCAG* |
| *Gapdh* | *F: GTGTCCGTCGTGGATCTGA*<br><br>*R: CCTGCTTCACCACCTTCTTG* |

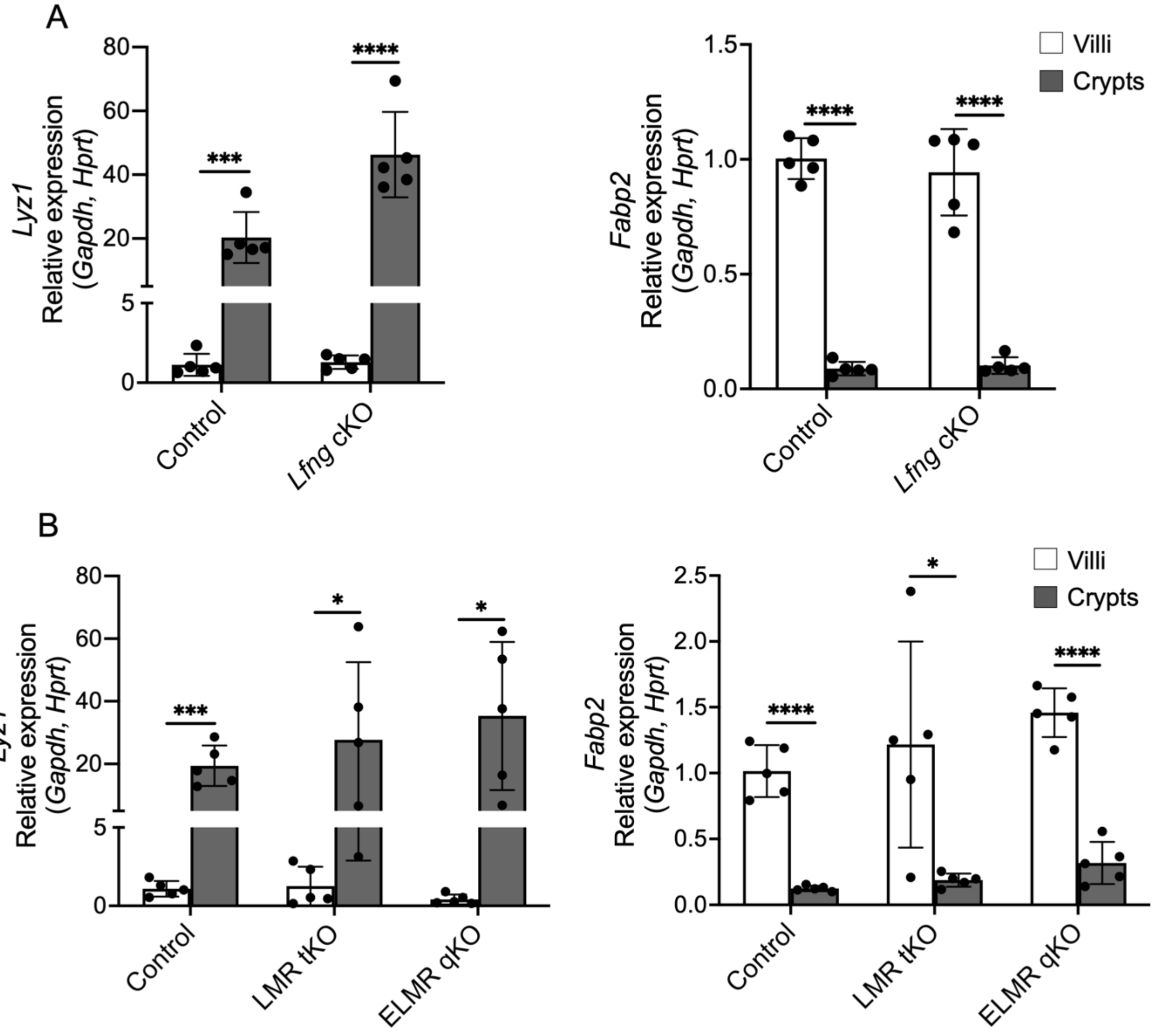


**Supplementary Figure S1**. Validation of villus and crypt fractions. (A) Enrichment of *Lyz1* in crypts (gray) and *Fabp2* in villi (white) is shown in Control and *Lfng* cKO mice (n = 5 mice per group). (B) Enrichment of *Lyz1* in crypts (gray) and *Fabp2* in villi (white) is shown Control, LMR tKO and ELMR qKO mice (n = 5 mice per group). *P* values from unpaired, two-tailed Student's t test – *$P < 0.05$, ***$P < 0.001$, ****$P < 0.0001$.

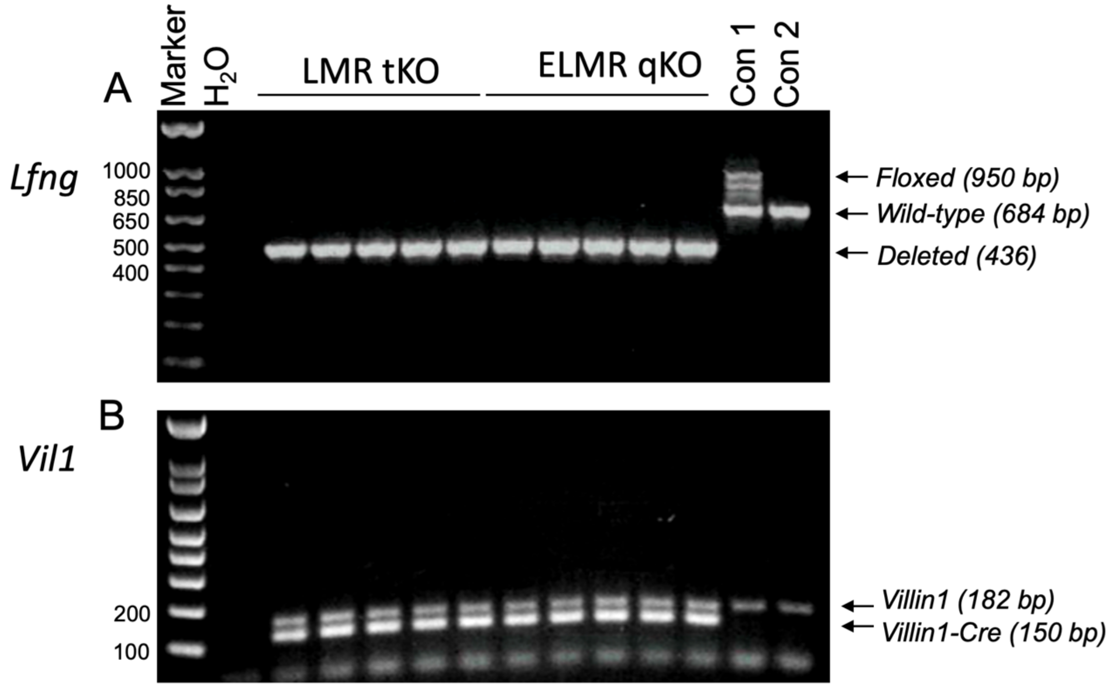


**Supplementary Figure S2.** Validation of *Lfng* deletion by *Villin1*-Cre in gDNA isolated from crypts of LMR tKO and ELMR qKO intestine. (A) *Lfng*[F] floxed (950 bp), *Lfng*[+] wild-type (684 bp) and *Lfng*[-] deleted bands (436 bp) (n = 5 mice per group). The *Vil1* endogenous gene (182 bp) and *Villin*-Cre transgene (~150 bp) were detected in the same mice (n = 5 mice per group). Con 1 and 2 were from *Lfng*[F/+] crypts lacking *Villin*-Cre.

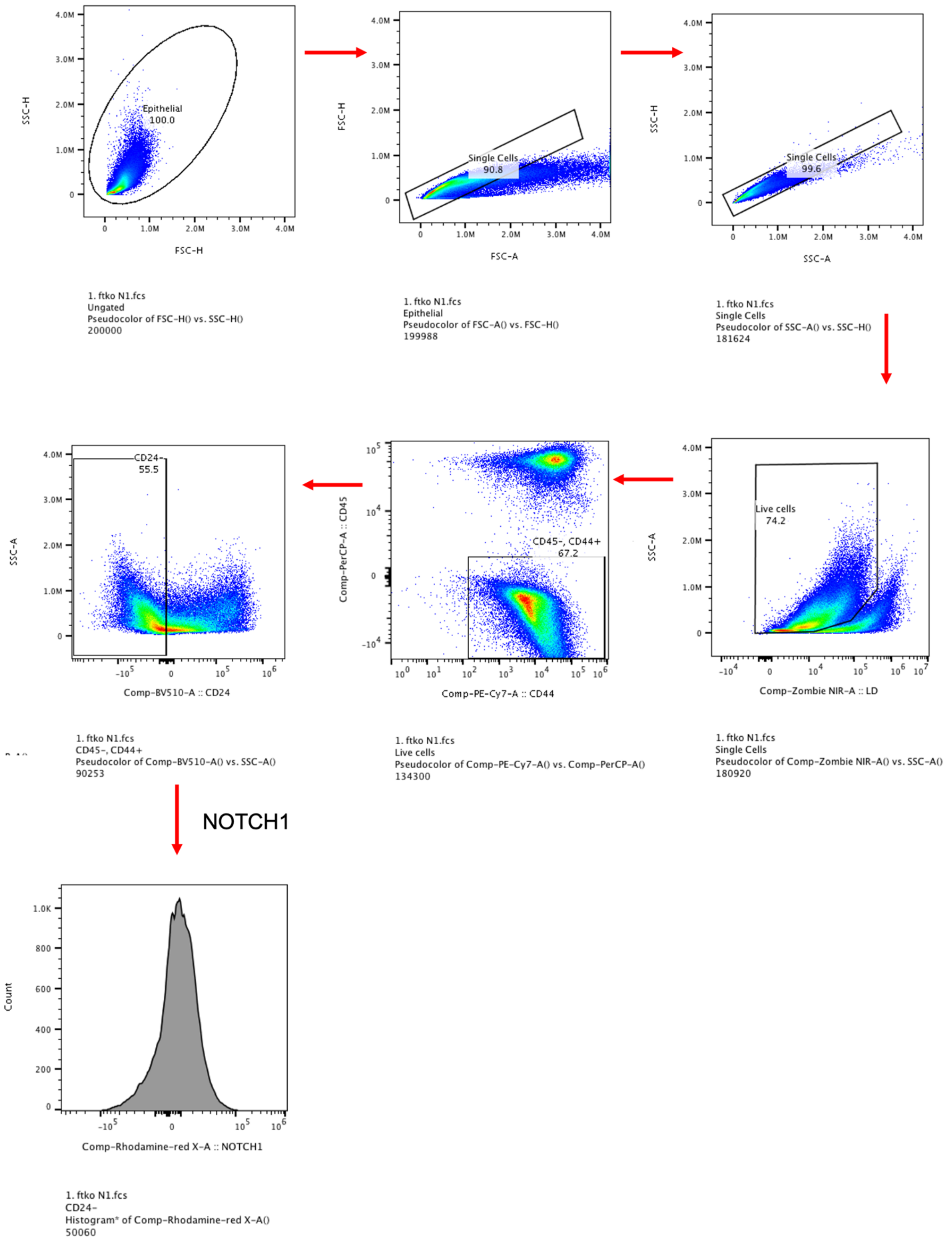
Epithelial
100.0
SSC-H
FSC-H
1. ftko N1.fcs
Ungated
Pseudocolor of FSC-H() vs. SSC-H()
200000
Single Cells
90.8
FSC-H
FSC-A
1. ftko N1.fcs
Epithelial
Pseudocolor of FSC-A() vs. FSC-H()
199988
Single Cells
99.6
SSC-H
SSC-A
1. ftko N1.fcs
Single Cells
Pseudocolor of SSC-A() vs. SSC-H()
181624
Live cells
74.2
SSC-A
Comp-Zombie NIR-A :: LD
1. ftko N1.fcs
Single Cells
Pseudocolor of Comp-Zombie NIR-A() vs. SSC-A()
180920
CD45-, CD44+
67.2
Comp-PerCP-A :: CD45
Comp-PE-Cy7-A :: CD44
1. ftko N1.fcs
Live cells
Pseudocolor of Comp-PE-Cy7-A() vs. Comp-PerCP-A()
134300
CD24-
55.5
SSC-A
Comp-BV510-A :: CD24
1. ftko N1.fcs
CD45-, CD44+
Pseudocolor of Comp-BV510-A() vs. SSC-A()
90253
NOTCH1
Count
Comp-Rhodamine-red X-A :: NOTCH1
1. ftko N1.fcs
CD24-
Histogram* of Comp-Rhodamine-red X-A()
50060

**Supplementary Figure S3.** Gating strategy for flow cytometry analysis. Single live cells were gated as shown to obtain the ISC cell population (CD45-CD44+CD24-) from single crypt cells. Mean fluorescence index (MFI) for anti-NOTCH1 ECD Ab binding or 2° Ab alone sample was determined on CD45-CD44+CD24- cells. The same gating strategy was used to determine DLL1-Fc or DLL4-Fc binding to ISC detected by anti-Fc-rhodamine Ab.
